# Targeting fibroblast activation suppresses chemotherapy-induced pro-invasive remodeling of the extracellular matrix

**DOI:** 10.64898/2026.09.04.749413

**Authors:** Anna Yui, Liyi Peng, Katie C. Lew, Dylan J. Landau, Crystal Zhang, Thomas J. Gerton, Marissa Li, Steffy Jullian Guyard, Mallory M. Caron, Jackson P. Fatherree, Rachel A. McGinn, Abraham L. Bayer, Pilar Alcaide, Charlotte Kuperwasser, Madeleine J. Oudin

## Abstract

Triple-negative breast cancer (TNBC) is an aggressive subtype of breast cancer that accounts for nearly 20% of breast cancer diagnoses. Despite recent approvals of novel therapeutic agents for TNBC, neoadjuvant chemotherapy remains part of standard of care treatment for TNBC patients. However, approximately 60% of TNBC patients who received neoadjuvant chemotherapy fail to achieve pathological complete response and face significantly elevated risks of recurrence and distant metastasis. We have shown that chemotherapy can induce significant changes in TNBC tumor extracellular matrix (ECM) that drive tumor cell invasion and may contribute to recurrence. Here, we investigate how chemotherapy drugs can impact resident fibroblasts, the major generators of ECM in the breast tissue. We find that chemotherapy activates fibroblasts and that doxorubicin specifically induces the production of pro-invasive ECM by fibroblasts via TGF-β/Smad3 signaling. An FDA-approved tyrosine kinase inhibitor nintedanib suppressed doxorubicin-induced fibroblast activation and pro-invasive ECM production by fibroblasts *in vitro*. Additionally, in a TNBC mouse model, nintedanib reduced fibroblast activation markers in tumors, suppressed tumor cell motility in the tumor ECM, and reduced lung metastasis. These findings demonstrate a novel mechanism by which chemotherapy can induce fibroblast activation and imply that targeting fibroblast activation reduces chemotherapy-induced metastasis in TNBC.

## Introduction

Breast cancer is a leading cause of cancer-related deaths in women. In the United States, approximately one in eight women will be diagnosed with breast cancer in their lifetime (1). Triple-negative breast cancer (TNBC) is an aggressive subtype of breast cancer, accounting for 15–20% of all breast cancer diagnoses. The particularly unfavorable prognosis of TNBC has historically been attributed to the absence of targetable receptors—estrogen receptor (ER), progesterone receptor (PR), and human epidermal growth factor receptor 2 (HER2)—which precludes the use of targeted therapies against these receptors. Although the recent approvals of the immune checkpoint inhibitor pembrolizumab (Keytruda) and the antibody-drug conjugates (ADCs) sacituzumab govitecan (Trodelvy) or datopotamab deruxtecan (Datroway) have expanded the treatment landscape (2–5), chemotherapy remains a central component of TNBC treatment in the neoadjuvant setting, both as a partner for combination regimens with immune checkpoint inhibitors and as the cytotoxic payload in ADCs (6). Moreover, given that the proportion of patients eligible for these newer treatment options remains limited, chemotherapy continues to be the most widely accessible treatment for TNBC patients (7,8). Even with aggressive chemotherapy regimens and new therapies, there is a high rate (up to 46%) of metastatic relapse (9–14) and the five-year survival for metastatic TNBC is 12%. Understanding the mechanisms that drive recurrence and chemotherapy resistance is critical to improve outcomes in TNBC.

Gene expression studies in TNBC after neoadjuvant chemotherapy reveal extracellular matrix (ECM) remodeling and wound healing gene signatures to be associated with poor responses and outcomes (15–18). The ECM is a major structural and signaling component of the tumor microenvironment (TME), providing both a physical scaffold for tissues and a reservoir for soluble growth factors and cytokines. In tumors, aberrant ECM remodeling, characterized by altered composition, stiffness, and organization, is well established as a driver of tumor progression and metastasis (19). We previously demonstrated that chemotherapy induces significant pro-invasive changes in the composition of the tumor ECM in an MMTV-PyMT TNBC mouse model and human TNBC samples (20). Within breast tissue, fibroblasts are the main cells that secrete and organize the ECM. In tumors, fibroblasts are converted into cancer-associated fibroblasts (CAFs) and can be classified into multiple subtypes based on their transcriptional profiles. Among them, myofibroblast-like CAFs (myCAFs), characterized by the expression of high levels of α-smooth muscle actin (α-SMA), drive tumor progression via ECM production and remodeling (21,22). On the other hand, inflammatory CAFs (iCAFs) are typically observed more distantly from tumor cells, express low levels of α-SMA and contribute to the induction of pro-tumorigenic TME through the secretion of inflammatory cytokines such as IL-6 and IL-11 (21). Additional subtypes such as vascular CAFs (vCAFs), characterized by the high expression of genes involved in vascularization and angiogenesis regulation, are identified through single-cell and spatial proteomics analysis of human breast tumors (23). Fibroblast activation is regulated by a complex network of signaling pathways, including but not limited to growth factors, inflammatory cytokines, and mechanotransduction signals (24). TGF-β- mediated Smad3 phosphorylation is recognized as a canonical driver of myCAF differentiation, promoting ECM production and the expression of α-SMA (25–27). IL-1-induced activation of Jak2/Stat3 pathway has been implicated in the induction of iCAFs, a subtype of CAFs known to secrete inflammatory cytokines such as IL-6 (21,25). Previous studies have shown an upregulation in α-SMA expression by immunostaining within the residual tumors of patient-derived xenograft model treated with doxorubicin and cyclophosphamide (28) and increase in ACTA2 transcript, the gene coding for α-SMA, in tumor single cells from human TNBC patients that did not respond to neoadjuvant chemotherapy compared to responders (29). However, whether chemotherapy can directly affect mammary fibroblasts and impact their ECM production or inflammatory function, and the mechanisms by which this occurs remain unknown.

In this study, we demonstrate that chemotherapy drugs activate normal mammary fibroblasts and that doxorubicin specifically induces changes in the ECM that drive tumor cell invasion and metastasis. We identify that the TGF-β/Smad3 pathway mediates chemotherapy-induced fibroblast activation and demonstrate that pharmacological inhibition of this pathway with an FDA-approved tyrosine kinase inhibitor nintedanib suppresses the pro-invasive changes in the fibroblast-derived ECM. We further show that the combination therapy of a syngeneic TNBC mouse model with doxorubicin and nintedanib suppresses doxorubicin-driven increase in lung metastasis without compromising the anti-tumor efficacy of doxorubicin or showing toxicity. Together, these findings uncover a mechanism of chemotherapy-induced fibroblast activation and provide a proof-of-concept for suppressing chemotherapy-induced pro-invasive changes in the ECM by the combination treatment with chemotherapy and an inhibitor of fibroblast activation.

## Results

### Chemotherapy is associated with increased **α**-SMA expression at the mRNA and protein levels

*ACTA2* is known to be highly expressed in myCAFs (21,23). We examined whether neoadjuvant chemotherapy exposure is associated with changes in *ACTA2* expression in primary breast tumor samples using publicly available datasets containing patient-matched primary breast tumor samples collected before and after neoadjuvant chemotherapy (30–34) (Table 1). Across all five datasets, between 41-78% of patients showed increased *ACTA2* expression following neoadjuvant chemotherapy. *ACTA2* expression was significantly increased following neoadjuvant chemotherapy in two datasets (Figure 1A, B, Fig. S1). These findings suggest that neoadjuvant chemotherapy is associated with increased *ACTA2* expression in primary breast tumors.

**Figure 1.**
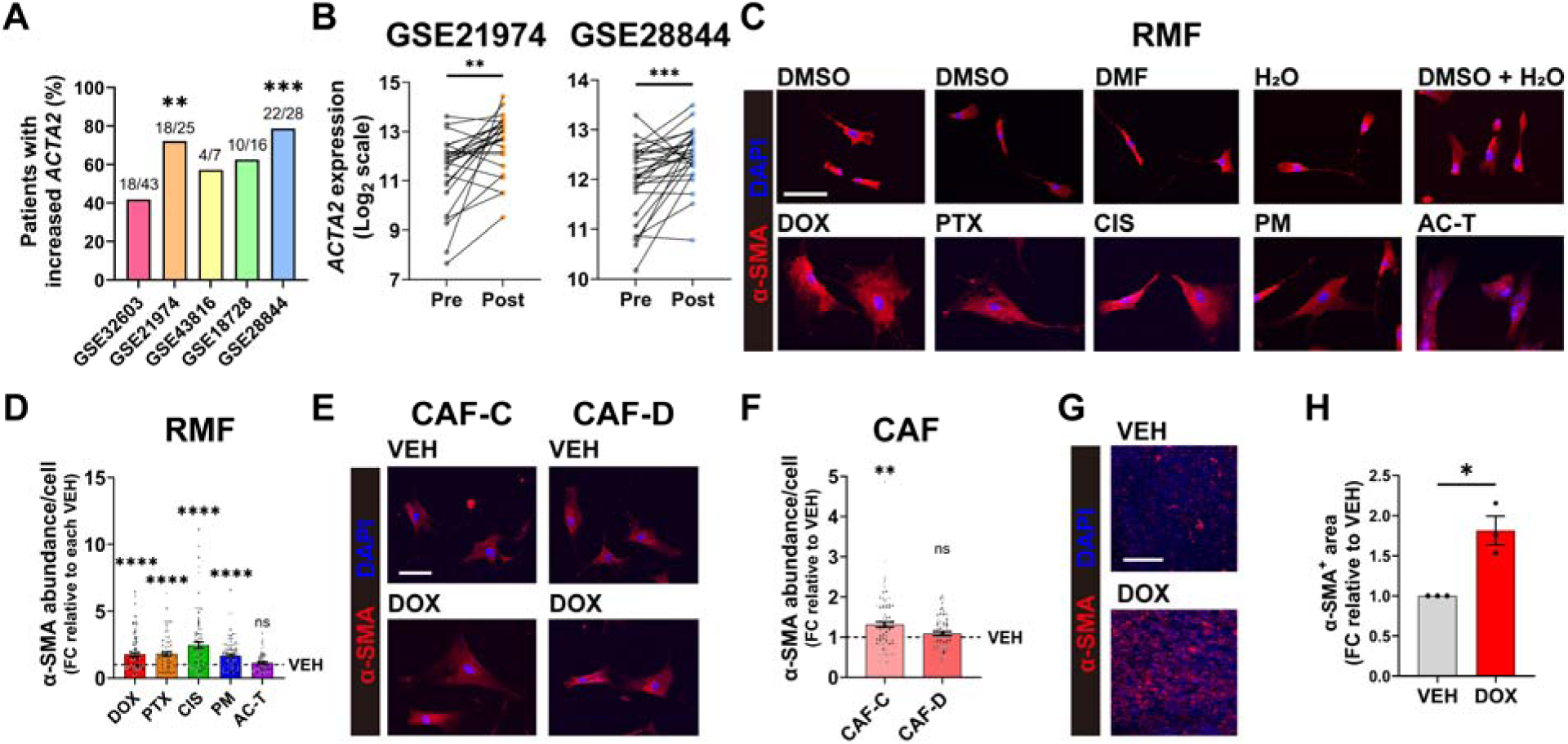
Chemotherapy increases CAF marker expression *in vivo* and *in vitro*. **A**) Analysis of *ACTA2* expression using microarray gene expression datasets from the GEO. The percentage of patients with increased *ACTA2* expression in post-chemotherapy samples is shown. **B)** *ACTA2* expression in patient-matched pre- and post-chemotherapy samples. **C**) Representative images of RMFs. For single-agent treatments, RMFs were treated with 100 nM doxorubicin, 5 nM paclitaxel, 5 µM cisplatin, 50 µM phosphoramide mustard, or vehicle control for 72 h. For AC-T treatment, RMFs were first treated with a mixture of 50 nM doxorubicin and 50 µM phosphoramide mustard or vehicle control for 72 h, followed by a 72 h of incubation in chemotherapy-free media, and subsequent treatment with 5 nM paclitaxel or vehicle control for 72 h. RMFs were immunostained for α-SMA (red), and DAPI (blue). Vehicle-treated cells of each condition are shown above the images of each drug-treated cells. Scale bar: 100 µm. **D**) Fold change (FC) in α-SMA integrated intensity, representing α-SMA abundance, was quantified. Data are shown as FC relative to the corresponding vehicle control (VEH). **E**) Representative images of CAFs treated with 100 nM doxorubicin or vehicle control for 72 h. CAFs were immunostained for α-SMA (red), and DAPI (blue). Scale bar: 100 µm. **F**) FC in α-SMA abundance was quantified. Data are shown as FC relative to the corresponding vehicle control. G) Representative images of tumors. EO771 tumor-bearing C57BL/6J mice were treated with doxorubicin (5 mg/kg) or vehicle control. Tumors were immunostained for α-SMA (red), and DAPI (blue). Scale bar: 100 µm. **H**) Area of α-SMA-positive areas in tumors were quantified. The tissues were immunostained for α-SMA (red) and DAPI (blue). Data show mean ± SEM [GSE32603: n=43; GSE21974: n=25; GSE43816: n=7; GSE18728: n=16; GSE28844: n=28 pairs of patient-matched pre- and post-neoadjuvant chemotherapy samples (**A, B**), n=3 biological replicates, at least 60 cells per condition in total (**C-F**), or n=3 independent mice per condition (**G, H**)]. Different-colored data points represent different biological replicates where applicable. Significance was determined by paired *t*-test (**A, B, H**) or unpaired *t*-test with Welch’s correction relative to their own vehicle (**D, F**). *p<0.05, **p<0.01, ***p<0.001, ****p<0.0001. DOX, doxorubicin; PTX, paclitaxel; CIS, cisplatin; PM, phosphoramide mustard; VEH, vehicle.

**Table 1.** ####

| Accession number | Platform | Pairs included in our analysis (n) | Treatment |
| --- | --- | --- | --- |
| GSE32603 (30) | GPL14668 | 43 | Anthracycline (AC) only; AC + taxane (T); AC + T + Trastuzumab; or AC + T + Other |
| GSE21974 (31) | GPL6480 | 25 | Epirubicine + cyclophosphamide followed by docetaxel |
| GSE43816 (32) | GPL11532 | 7 | Epirubicine + cyclophosphamide followed by docetaxel |
| GSE18728 (33) | GPL570 | 16 | Docetaxel and capecitabine (TX); or TX followed by adriamycin + cyclophosphamide |
| GSE28844 (34) | GPL570 | 28 | Epirubicin + cyclophosphamide followed by paclitaxel + gemcitabine ± trastuzumab; doxorubicin + pemetrexed followed by docetaxel; or doxorubicin + cyclophosphamide followed by docetaxel |

Chemotherapy treatment affects both normal mammary fibroblasts and CAFs (24). We next evaluated the effects of individual chemotherapy drugs on the activation of reduction mammary fibroblasts (RMFs), derived from healthy human breast tissue (35). We treated RMFs individually with each clinically used chemotherapy drugs (doxorubicin, paclitaxel, cisplatin, and cyclophosphamide) or in combination as the AC-T regimen (doxorubicin and cyclophosphamide followed by paclitaxel). Since cyclophosphamide is a prodrug and inactive in cell culture, a reactive metabolite phosphoramide mustard was used for *in vitro* work. Doxorubicin, cisplatin, and cyclophosphamide are DNA-damaging agents, and paclitaxel is an antimicrotubule agent (36–39). Drug concentrations were selected to achieve a 30–50% reduction in RMF number (Fig. S2A). These drug concentrations induced moderate cytotoxicity in a human TNBC cell line MDA-MB-231, while drug sensitivity varied among additional human TNBC cell lines (Sup. Table 1). First, RMFs treated with individual drugs or AC-T exhibited significantly greater cell area than their respective vehicle controls (Fig. S2B, C). Further, all individual drugs showed significantly higher α-SMA abundance compared to each vehicle control (Figure 1C, D). We then investigated whether two breast cancer patient-derived lines (CAF-C and CAF-D) (40) responded similarly, focusing on doxorubicin, which we have shown in mouse models has the strongest effects on ECM composition and abundance in mouse tumors(20). Doxorubicin increased α-SMA abundance in CAF-C relative to vehicle control, whereas no significant difference was observed in CAF-D (Figure 1E, F). To investigate if chemotherapy increases CAF markers in mice, we used the EO771 syngeneic TNBC mouse model treated with doxorubicin or vehicle control. Expression area of α-SMA was significantly increased in tumors of doxorubicin-treated mice (Figure 1G, H), indicating enhanced fibroblast activation after doxorubicin treatment. Together, these results demonstrate that *ACTA2* expression is elevated in some portions of breast cancer patients treated with neoadjuvant chemotherapy. Further, we find that chemotherapy drugs can elevate α-SMA expression and induce pro-fibrotic myCAF-like phenotype (21,22) in normal mammary fibroblasts.

### Doxorubicin and cisplatin activate RMFs via TGF-**β**/Smad3 pathway and doxorubicin uniquely induces pro-invasive changes in fibroblast-derived ECM

We next investigated the mechanisms by which chemotherapy affects fibroblasts to drive increased activation. We first examined whether chemotherapy drugs impact TGF-β-mediated Smad3 phosphorylation, which is recognized as a canonical driver of myCAF differentiation, and promotes ECM production and the expression of α-SMA (22,25,41). Doxorubicin and cisplatin significantly upregulated Smad3 phosphorylation at Ser423/425 in RMFs while paclitaxel significantly downregulated it (Figure 2A, B). Phosphoramide mustard and AC-T had no significant effect on Smad3 phosphorylation.

**Figure 2.**
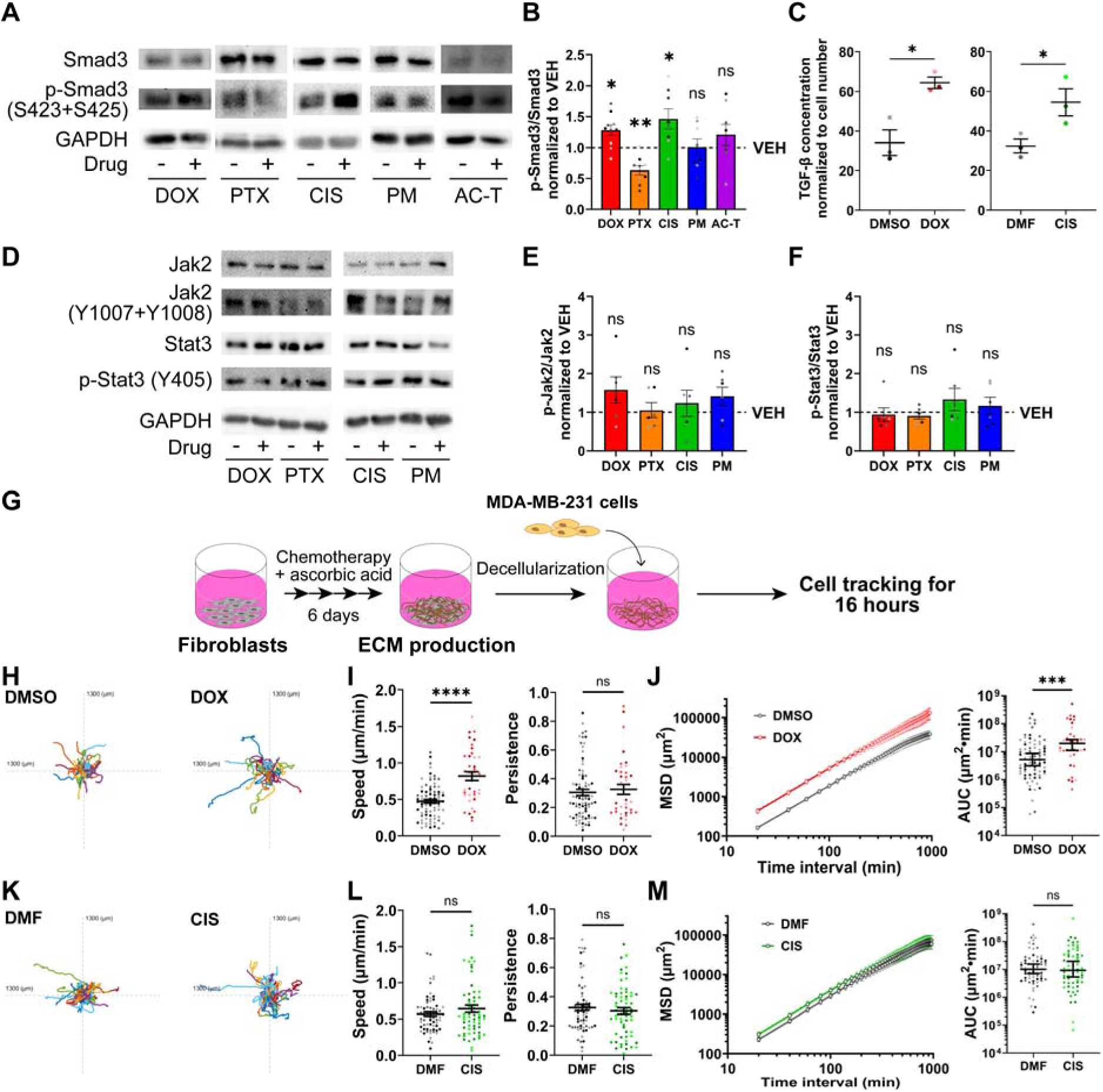
**Doxorubicin and cisplatin upregulate TGF-**β**/Smad3 pathway. A, D**) Representative blots of whole cell lysates from RMFs treated with chemotherapy. For single-agent treatments, RMFs were treated with 100 nM doxorubicin, 5 nM paclitaxel, 5 µM cisplatin, 50 µM phosphoramide mustard, or vehicle control for 72 h. For AC-T treatment, RMFs were first treated with a mixture of 50 nM doxorubicin and 50 µM phosphoramide mustard or vehicle control for 72 h, followed by a 72 h of incubation in chemotherapy-free media, and subsequent treatment with 5 nM paclitaxel or vehicle control for 72 h. **B**) Quantification of phospho-Smad3 (Ser423 + Ser425) protein relative to total Smad3 protein. Band intensities were normalized to a loading control (GAPDH). **C**) Human TGF-β1 ELISA using conditioned media of RMFs treated with 100 nM doxorubicin, 5 µM cisplatin, or vehicle control for 72 h. TGF-β1 concentrations were normalized to cell number. Doxorubicin and cisplatin significantly increased TGF-β1 secretion per cell. **E, F**) Quantification of phospho-Jak2 (Tyr1007 + Tyr1008) protein and phospho-Stat3 (Tyr705) relative to total Jak2 and Stat3 protein, respectively. Band intensities were normalized to a loading control (GAPDH). **G**) Schematic of reseeding experiments using fibroblast-derived ECM. Treatment schedules varied depending on the chemotherapy regimen. For all treatment regimens, RMFs were treated with the same drug concentrations as in **A**) or with vehicle control, and culture media were supplemented with 75 μg/mL ascorbic acid throughout the treatment period. For single-agent treatments, RMFs were treated with a drug for 6 days. 231-GFP+ cells were reseeded after the decellularization, incubated for 6 h, and imaged for 16 h. **H, K**) Trajectories of 231-GFP cells in RMF-derived ECM. The x- and y-axes range from −1300 to +1300 μm. **I, L**) Migration speed and persistence of 231-GFP cells. **J, M**) Mean squared displacement (MSD) curves and MSD AUC of 231-GFP cells. RMFs were treated with the same concentration of drugs indicated in the legend of Figure 1. Data show mean ± SEM (**B, C, E, F, I, J-MSD curves, L, M-MSD curves**) or median ± 95% CI (**J-AUC, M-AUC**) [n=4-5 independent cell lysates (**A, B**), n=3 independent conditioned media (**C**), n=3 independent cell lysates (**D-F**), or n=3 biological replicates, at least 37 cells per condition in total (**H-M**)]. Different-colored data points represent different biological replicates. Significance was determined by paired *t*-test (**B, C, E, F**), unpaired *t*-test with Welch’s correction (**I, L**), or Mann-Whitney test (**J, M**). Each drug was compared with its corresponding vehicle control. *p<0.05, **p<0.01, ***p<0.001, ****p<0.0001. DOX, doxorubicin; PTX, paclitaxel; CIS, cisplatin; PM, phosphoramide mustard; VEH, vehicle.

Doxorubicin and cisplatin significantly enhanced TGF-β1 secretion per cell, as measured in the conditioned media of drug-treated RMFs (Figure 2C). IL-1-mediated activation of Jak2/Stat3 pathway has also been implicated in the induction of iCAFs, a subtype of CAFs known to secrete inflammatory cytokines (25). None of the drugs significantly affected the phosphorylation of either Jak2 or Stat3 (Figure 2D-F). These data suggest that doxorubicin and cisplatin drive RMFs to a myCAF-like phenotype by activating TGF-β/Smad3 signaling, while paclitaxel, phosphoramide mustard, and AC-T may activate RMFs through different pathways.

To determine whether chemotherapy-induced fibroblast activation and downstream Smad3 phosphorylation drive changes in ECM production that can impact tumor cells, RMFs were treated with chemotherapy and allowed to produce ECM, which was subsequently decellularized to generate ECM scaffolds. GFP-labeled MDA-MB-231 TNBC cells (231-GFP) were reseeded into the scaffolds, and cell motility was tracked for 16 hours (Figure 2G). In ECM derived from doxorubicin-treated RMFs, 231-GFP cells invaded significantly faster, with no changes in persistence (Figure 2H, I). 231-GFP cells reseeded in ECM from doxorubicin-treated fibroblasts exhibited higher cell motility throughout all time intervals, as measured by the mean squared displacement (MSD), supporting enhanced cell motility over both short and long timescales (Figure 2J). To comprehensively evaluate the temporal dynamics of cell migration, we calculated the area under the curve (AUC) of the MSD, which provides a robust measure of net translocation that integrates both speed and directionality while minimizing sensitivity to tracking noise (42). Our data showed that MSD AUC was significantly higher in doxorubicin (Figure 2J). Overall, these data show that doxorubicin-treated ECM drives tumor cell invasion. In contrast, paclitaxel, cisplatin and phosphoramide mustard treatment of RMFs did not induce significant changes in speed and persistence, and paclitaxel induced a significant decrease in MSD AUC of 231-GFP cells (Figure 2K-M and Fig. S3A- H). The combination regimen AC-T led to a significant increase in cell invasion, a decrease in persistence and an increase in the AUC of the MSD (Fig. S3I-L).

Although doxorubicin and cisplatin both upregulate TGF-β/smad3 pathway in RMFs, only doxorubicin induces pro-invasive changes in fibroblast-derived ECM. Further, while AC-T did not upregulate TGF- β/Smad3 signaling, it did significantly increase the pro-invasive effects of the ECM. Together, these data indicate that doxorubicin uniquely drives TGF-β/Smad3 signaling in fibroblasts and induces pro-invasive changes in fibroblast-derived ECM.

### Inhibition of TGF-**β**/Smad3 pathway suppresses doxorubicin-induced fibroblast activation and reduces pro-invasive effects of fibroblast-derived ECM

To further dissect the roles of TGF-β/Smad3 pathway in driving doxorubicin-induced fibroblast activation and ECM production, we used different inhibitors of the canonical TGF-β/Smad3 pathway: Nintedanib, a tyrosine kinase inhibitor which targets PDGFR, FGFR, and VEGFR that has been shown to inhibit the TGF-β/Smad3 pathway and is FDA-approved for idiopathic pulmonary fibrosis (IPF) (43–45), galunisertib, which is a selective inhibitor of TGF-β receptor I (46) and 1D11 is anti-pan-TGF-β antibody (47). RMFs were treated with doxorubicin alone, doxorubicin in combination with the indicated inhibitor, or the corresponding vehicle controls. All inhibitors suppressed the doxorubicin-induced increase of α- SMA abundance in RMFs (Figure 3A, B). Combination of doxorubicin and nintedanib also resulted in a significantly smaller fibroblast cell area than doxorubicin alone (Fig. S4A, B). We then used SB202190 a pharmacologic inhibitor of p38, which mediates non-canonical TGF-β pathway signaling. In addition, we used Dickkopf-1 (Dkk-1), a Wnt signaling antagonist that is downregulated by TGF-β signaling that has been shown to reduce TGF-β-induced fibrosis when overexpressed *in vivo* (48). SB202190 but not Dkk-1 suppressed doxorubicin-induced increase of α-SMA abundance in RMFs (Figure 3A, B). We then tested the inhibitory effect of nintedanib and galunisertib on TGF-β/Smad3 pathway in RMFs. Consistent with previous observations, treatment with doxorubicin alone increased Smad3 phosphorylation. Notably, both nintedanib and galunisertib reduced Smad3 phosphorylation in doxorubicin-treated RMFs (Figure 3C). These data indicate that activation of the TGF-β/Smad3 pathway is essential for doxorubicin-driven fibroblast activation. We then investigated whether nintedanib affects doxorubicin-induced effects on CAFs (Figure 3D). The combination treatment with doxorubicin and nintedanib significantly reduced α- SMA abundance relative to doxorubicin alone (Figure 3E, F). These results suggest that the combination of doxorubicin and nintedanib could potentially not only suppress conversion of normal fibroblasts into CAFs but also reverse activated fibroblasts.

**Figure 3.**
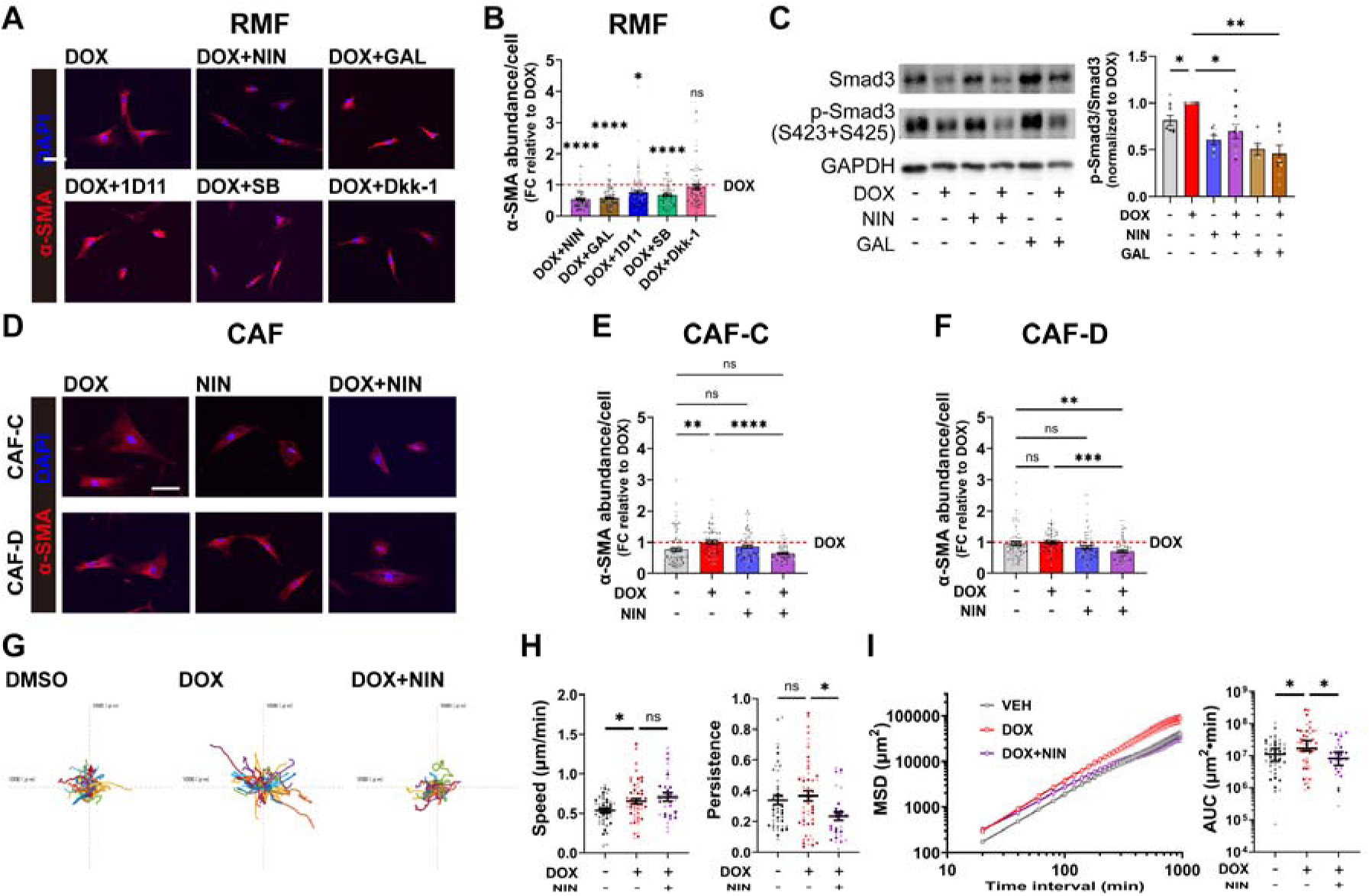
**TGF-**β**/Smad3 pathway inhibition suppresses doxorubicin-induced activation of fibroblasts and pro-invasive changes in ECM. A**) Representative images of RMFs treated with 100 nM doxorubicin, or 100 nM doxorubicin and the indicated inhibitor (1 μM nintedanib, 5 μM galunisertib, 10 μg/mL 1D11 antibody, 20 μM SB202190, or 100 ng/mL Dkk-1) for 72 h. The same concentrations were used when applicable throughout the experiments shown in this figure. RMFs were immunostained for α-SMA (red), and DAPI (blue). **B**) Fold change (FC) in α-SMA abundance was quantified. **C**) Representative blots of whole cell lysates of RMFs treated with indicated conditions for 72 h and quantification of phospho-Smad3 (Ser423 + Ser425) protein relative to total Smad3 protein. Band intensities were normalized to a loading control (GAPDH). Data are shown as FC to doxorubicin. **D**) Representative images of CAF-C or CAF-D treated with doxorubicin alone, nintedanib alone, or doxorubicin and nintedanib for 72 h. CAFs were immunostained for α-SMA (red), and DAPI (blue). **E, F**) FC in α-SMA abundance was quantified. **G**) Trajectories of 231-GFP cells in RMF-derived ECM. The x- and y-axes range from −1000 to +1000 μm. RMFs were treated with the indicated conditions with 75 μg/mL ascorbic acid for 6 days. **H**) Migration speed and persistence of 231-GFP cells. **I**) MSD curves and MSD AUC of 231-GFP cells. Data show mean ± SEM (**B, C, E, F, H, I-MSD curves**) or median ± 95% CI (**I-AUC**) [n=3 biological replicates, at least 71 cells per condition in total (**A, B, D-F**), n=3-5 independent cell lysates (**C**), or n=3 biological replicates, at least 30 cells per condition in total (**G-I**)]. Different-colored data points represent different biological replicates. Significance was determined by ordinary one-way ANOVA followed by Dunnett’s multiple comparisons test (**B, H**) or Šídák’s multiple comparisons test (**E, F**), mixed-effects analysis followed by Šídák’s multiple comparisons test (**C**), or Kruskal-Wallis test followed by Dunn’s multiple comparisons test (**I**). Doxorubicin was compared with the corresponding vehicle control or the combination of doxorubicin and the indicated inhibitor. *p<0.05, **p<0.01, ***p<0.001, ****p<0.0001. DOX, doxorubicin; NIN, nintedanib; GAL, galunisertib; SB, SB202190.

To assess whether the inhibition of TGF-β/Smad3 signaling attenuates doxorubicin-induced pro-invasive ECM remodeling, we reseeded tumor cells onto decellularized ECM scaffolds derived from fibroblasts that received combination treatments to evaluate effects on cell motility. Treatment of RMFs with galunisertib, 1D11, or SB202190 did not significantly change migration speed or persistence of the 231- GFP cells relative to doxorubicin alone (Fig. S4C-G). Treatment of RMFs with nintedanib did not reduce cell invasion speed, but the combination of doxorubicin and nintedanib exhibited significantly reduced persistence relative to doxorubicin (Figure 3G, H and Fig. S4F, G). While the MSD curve of doxorubicin revealed increased cell motility across all time intervals relative to vehicle control, the MSD curve for the doxorubicin + nintedanib overlapped with that of doxorubicin at short time intervals but progressively converged to vehicle control’s curve at longer time intervals (Figure 3I). Quantification of MSD AUC showed that doxorubicin has significantly higher value than vehicle control and doxorubicin + nintedanib, indicating that doxorubicin-induced increase in cell motility can be suppressed by nintedanib, but not by the other inhibitors tested (Figure 3I and Fig. S4H). This pattern indicates that, despite maintaining comparable motility, since 231-GFP cells on ECM derived from doxorubicin + nintedanib-treated RMFs exhibit reduced directionality, it resulted in diminished net displacement. This highlights a decoupling of migration speed and effective long-range displacement in doxorubicin + nintedanib-treated RMFs’ ECM. Together, these data demonstrate that inhibition of the TGF-β/Smad3 pathway suppresses doxorubicin- induced fibroblast activation as indicated by reduced α-SMA and Smad3 phosphorylation, whereas only an FDA-approved tyrosine kinase inhibitor nintedanib suppressed pro-invasive effects of fibroblast- derived ECM. Given the established tyrosine kinase inhibitory activity of nintedanib (44), these findings suggest that its effects on the ECM may involve both tyrosine kinases and TGF-β/Smad3 pathway inhibition.

### Nintedanib reduces doxorubicin-induced metastasis in TNBC mouse model

We next aimed to investigate the potential of targeting chemotherapy-induced fibroblast activation to improve outcomes for TNBC. We used the FDA-approved drug nintedanib in combination with doxorubicin in an immunocompetent TNBC mouse model. We generated a syngeneic TNBC mouse model by injecting EO771 mouse TNBC cells into the fourth mammary fat pad of C57BL/6J mice, and determined the effect of nintedanib treatment in combination with doxorubicin. Mice in all treatment groups maintained stable body weight and body condition scores throughout the study, with no apparent signs of treatment-related toxicity (Fig. S5A). Doxorubicin significantly reduced endpoint tumor volume and there was no significant difference from the doxorubicin + nintedanib group (Figure 4A, B). However, doxorubicin led to a significant increase in lung metastasis, which was significantly reduced by the addition of nintedanib, suggesting that the combination of doxorubicin and nintedanib could potentially reduce the doxorubicin-induced increase in lung metastasis (Figure 4C, D, Fig. S5B).

**Figure 4.**
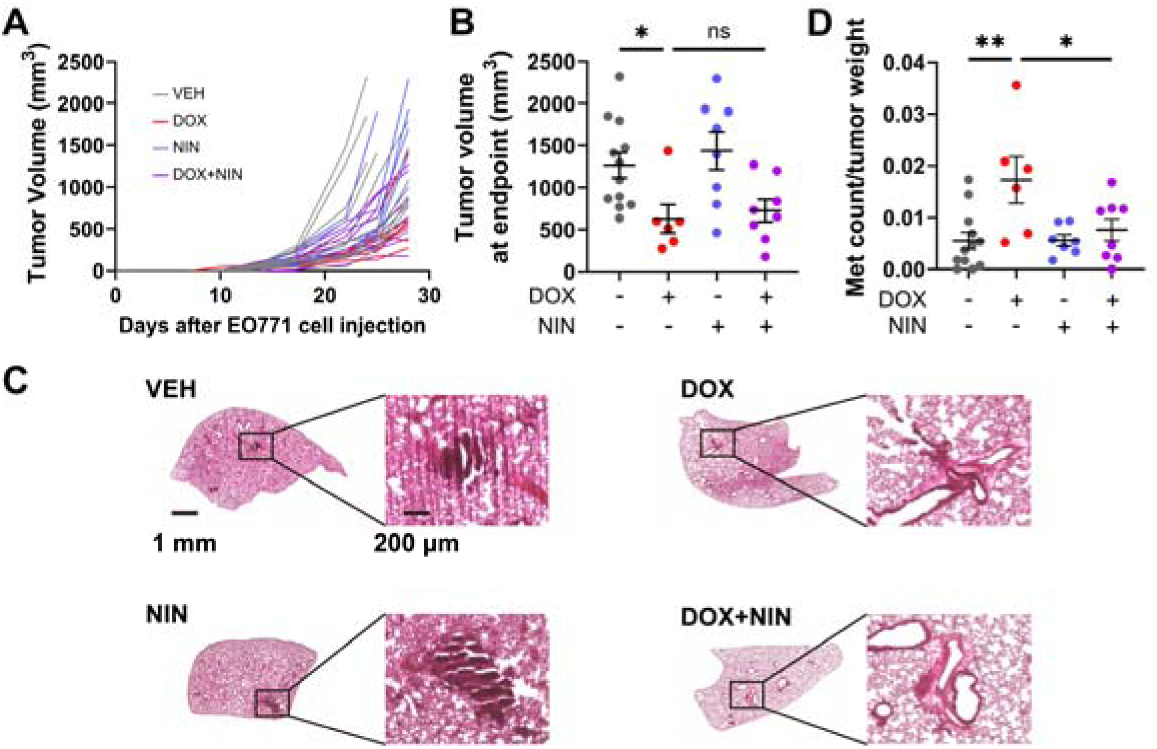
Nintedanib attenuates doxorubicin-induced lung metastatic efficiency without compromising doxorubicin’s anti-tumor efficacy. EO771 tumor-bearing C57BL/6J mice were treated with doxorubicin (5 mg/kg), nintedanib (50 mg/kg), doxorubicin with nintedanib, or corresponding vehicle controls. **A**) Individual tumor growth curve. Tumor size was measured using a digital caliper twice a week. Tumor volume was calculated using the equation “volume = 0.5 x length x width x width”, where length is the longest axis and width is the axis perpendicular to the length. **B**) Tumor volume at endpoint. **C**) Representative images of H&E-stained lungs. **D**) The number of metastatic foci was divided by tumor weight. Data show mean ± SEM [n=12, 6, 8, and 8 mice for vehicle, doxorubicin, nintedanib, and doxorubicin + nintedanib, respectively. For **C** and **D**, nintedanib group included 7 mice.]. Significance was determined by Kruskal-Wallis test followed by Dunn’s multiple comparisons test (**B**) or ordinary one-way ANOVA followed by Šídák’s multiple comparison test (**D**). *p<0.05, **p<0.01, ***p<0.001, ****p<0.0001. DOX, doxorubicin; NIN, nintedanib; VEH, vehicle.

### Doxorubicin + nintedanib suppresses CAF marker expression in TNBC tumors

We next investigated whether nintedanib treatment suppresses doxorubicin-induced fibroblast activation in the tumors. The tumors were immunostained for α-SMA as well as fibroblast activation protein (FAP), which is known to be highly expressed in CAFs involved in ECM remodeling (49,50) (Figure 5A).

**Figure 5.**
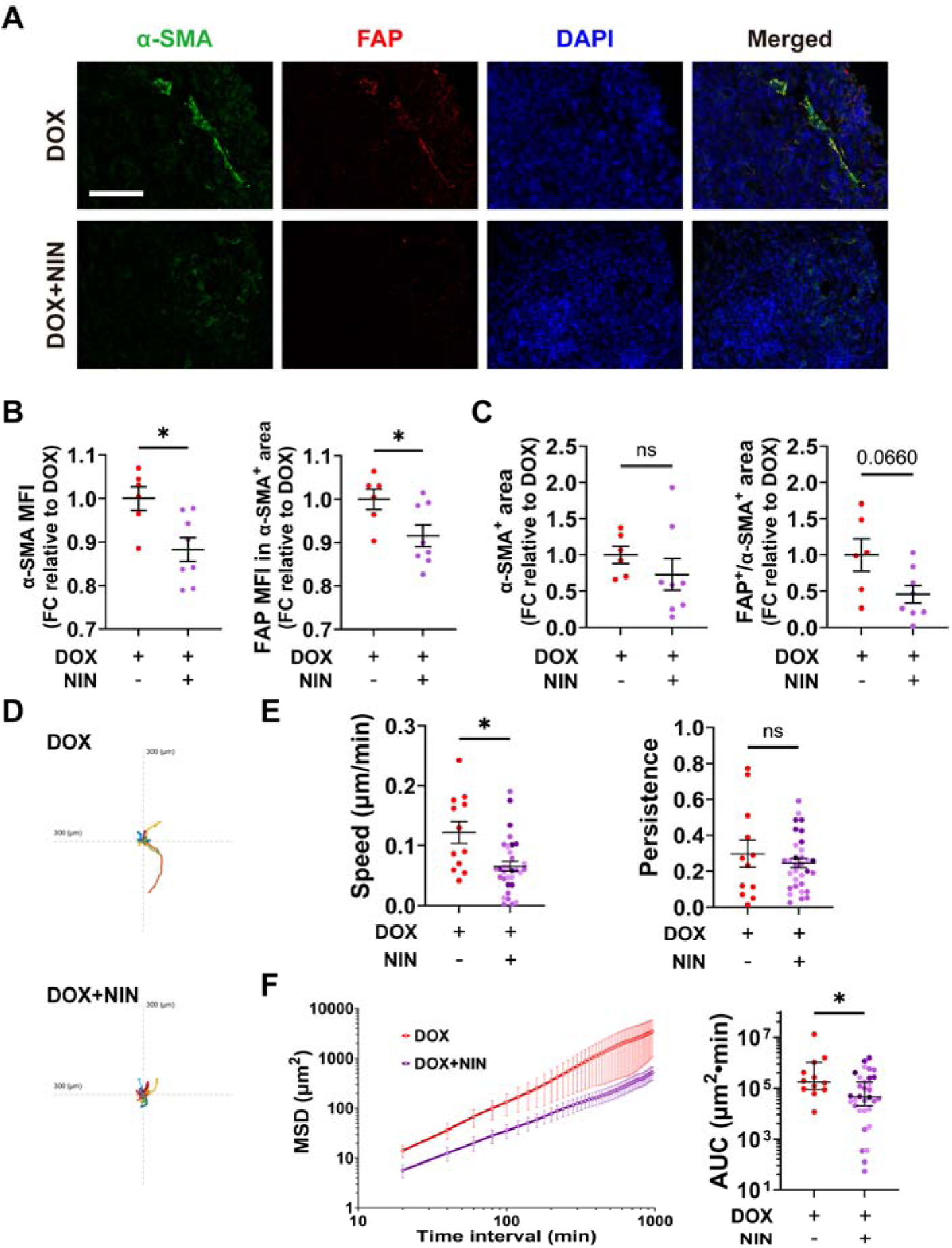
Nintedanib suppresses CAF markers in mouse TNBC tumors. **A**) Representative images of tumors. The tumors were immunostained for α-SMA (green), FAP (red), and DAPI (blue). Scale bar: 100 µm. **B**) MFI of α-SMA (left) and FAP in FAP-/α-SMA-positive regions (right) were quantified. **C**) Total area of α-SMA-positive regions and FAP-/α-SMA-positive regions were quantified. Quantified values were normalized to the mean value of doxorubicin-treated tissues stained and imaged on the same day to minimize inter-batch variability (**B, C**). Each data point represents a single mouse. **D**) Trajectories of 231- GFP cells in decellularized tumors. The x- and y-axes range from −300 to +300 μm. **E**) Migration speed and persistence of 231-GFP cells. **F**) MSD curves of 231-GFP cells and AUC calculated based on the MSD curves of individual cells. Data show mean ± SEM (**B, C, E, F-MSD curves**) or median ± 95% CI (**F-AUC**) [Doxorubicin: n=6, doxorubicin + nintedanib: n=8 (**A-C**), or doxorubicin: n=1, doxorubicin + nintedanib: n=4, at least 12 cells per condition in total (**D-F**).]. Significance was determined by unpaired *t*-test with Welch’s correction (**B, C, E**) or Mann-Whitney test (**F**). *p<0.05 was considered statistically significant. DOX, doxorubicin; NIN, nintedanib.

Treatment with doxorubicin and nintedanib decreased the mean fluorescence intensity (MFI) of α-SMA in tumors. We then quantified the MFI of FAP in the area with positive α-SMA staining to account for FAP expression in myCAFs, since FAP expression is also observed in tumor-associated macrophages in breast stroma (51). Here, we also observed a decrease in α-SMA/FAP positive cells in doxorubicin + nintedanib- treated tumors, suggesting the decrease of fibroblasts involved in ECM remodeling after the combination treatment (Figure 5B). Although we did not observe a significant decrease of α-SMA-positive area and FAP-/α-SMA-positive area, the decrease in FAP-/α-SMA-positive area showed trend toward decrease (p=0.0660) (Figure 5C).

Lastly, to investigate the effects of the combination treatment on cancer cell invasion, the tumors were decellularized to obtain ECM scaffolds as described previously (52). 231-GFP cells were reseeded onto the scaffolds, and cell motility was tracked overnight. We observed a significant decrease in the invasion speed of the cells reseeded onto the scaffolds derived from doxorubicin + nintedanib-treated mice compared to the cells reseeded onto doxorubicin-treated mice’s scaffolds, with no effects on persistence (Figure 5D, E). Doxorubicin + nintedanib consistently exhibited lower MSD than doxorubicin alone across all time intervals and had a significantly lower MSD AUC (Figure 5F). Together, these data demonstrate that nintedanib reduces fibroblast activation and decreases the pro-invasive effects of the ECM in doxorubicin-treated tumors.

## Discussion

Chemotherapy treatment remains a pillar of cancer treatment, particularly for TNBC, even with the addition of newer modalities such as immunotherapy and ADCs. Given the high rate of metastatic relapse (up to 46%) (9–14), it is critical that we increase our understanding of how chemotherapy treatment could be contributing to dissemination and ultimately relapse. Chemotherapy drugs, whether delivered systemically or locally, do not only target tumor cells, but also can impact cells in the stroma. Our previous work had shown that chemotherapy can induce distinct pro-invasive changes in the ECM of TNBC tumors (20), however, the mechanism by which this occurs was unknown. Here, we show that chemotherapy treatment can increase the activation of fibroblasts as measured by α-SMA in patients, mouse models and patient-derived fibroblasts and CAFs. Doxorubicin activates fibroblasts by upregulating TGF-β/Smad3 pathway, which resulted in enhanced motility of MDA-MB-231 cells within the ECM produced by fibroblasts, effects that were suppressed by tyrosine kinase inhibitor nintedanib. In a TNBC mouse model, nintedanib decreased the expression of CAF markers and MDA-MB-231 cells’ motility in the tumors and suppressed lung metastasis. Together, these data provide the first mechanism by which chemotherapy, specifically doxorubicin, induces pro-invasive changes in ECM production by breast tissue fibroblasts that contribute to metastatic progression.

Our results bring attention to the unintended effects that systemic non-targeted treatments have on stromal cells within the primary tumor. All tested chemotherapy drugs individually and our *in vitro* version of the AC-T TNBC regimen increased α-SMA expression levels in breast fibroblasts. However, only the DNA-damaging drugs doxorubicin and cisplatin significantly increased phosphorylation of Smad3, and only doxorubicin induced pro-invasive changes in the ECM produced by RMFs. Paclitaxel and phosphoramide mustard had no effect on Smad3 or Jak/Stat signaling, and did not impact ECM-driven invasion. This is in line with recent work which demonstrated drug-specific effects on CAFs, showing that doxorubicin induces apoptosis in certain populations of ECM-producing myCAFs, whereas paclitaxel primarily alters their motility and morphology with minimal impact on cell survival (53). Another study found that chemotherapy drugs such as carboplatin, doxorubicin, and paclitaxel, can induce a catabolic CAF-like phenotype *in vitro* and stimulate antioxidant and immune signaling pathways in human foreskin fibroblasts that support tumorigenic properties in co-cultured MCF7 cells (54). While the distinct mechanisms by which chemotherapy drugs impact fibroblast function are not identified in these studies, these findings further support drug-specific effect on ECM production. Both doxorubicin and platinum-based drugs like cisplatin and carboplatin function as DNA-damaging agents, although via different mechanisms: Doxorubicin triggers DNA double-strand breaks by poisoning topoisomerase II following its intercalation into the DNA helix (36), while cisplatin forms stable platinum-DNA adducts, primarily in the form of intrastrand cross-links (38). Paclitaxel induces cell death via microtubule stabilization (37) and cyclophosphamide undergoes metabolic activation and exhibits cytotoxicity by alkylating guanine residues in DNA, resulting in intra- and interstrand DNA crosslinks (39), suggesting that different mechanisms are involved in mediating chemotherapy-driven effects on fibroblast phenotypes in the primary tumor. In addition to chemotherapy, adjuvant radiation therapy remains an integral component of the standard of care for TNBC, particularly for patients with locally advanced disease or those undergoing breast-conserving surgery (55). Irradiated fibroblasts adopt a unique secretory profile through metabolic adaptations, creating a pro-tumorigenic microenvironment that promotes the progression of TNBC (56). These findings all support that importance of understanding how different chemotherapy drugs and radiation, individually and in combination, impact fibroblasts and their pro-tumorigenic properties, to understand the development of drug resistance and recurrence.

In tumors, fibroblasts differentiate into heterogenous populations of cancer-associated fibroblasts (CAFs) which exhibit different transcriptional profiles that dictate specialized functions and impose different impacts on tumor cells (57). Here, we focus on the effects of chemotherapy drugs on ECM production in tumors, which is largely driven by myCAFs (22). Several markers of myCAFs have been reported, with α-SMA expression is a crucial determinant of fibroblast contractile activity, driving the mechanical remodeling and contraction of the ECM (58). Further, different pathways that drive ECM production have been identified in myCAFs (25,59–63). In RMFs, both doxorubicin and cisplatin increase α-SMA, TGF-β secretion and activation of the Smad3 pathway signaling, however only doxorubicin leads to pro-invasive changes in ECM production, suggesting that neither chemotherapy-induced increase of α-SMA nor its concurrent increase together with TGF-β/Smad3 pathway upregulation is sufficient to induce pro-invasive changes in the ECM produced by normal mammary fibroblasts. Pharmacological inhibition of canonical and noncanonical TGF-β signaling components suppressed doxorubicin-induced α-SMA expression levels in RMFs suggesting the involvement of TGF-β receptors in this process. Nintedanib, which suppresses phosphorylation of PDGFR, FGFR, and VEGFR in primary human lung fibroblasts derived from IPF patients (64), as well as Smad3 in normal human fetal diploid fibroblasts (45), was the only inhibitor that was successful in reducing doxorubicin-induced pro-invasive effects in this study. Galunisertib has been reported to significantly attenuate TGF-β-induced phosphorylation of Smad2/3 in human dermal fibroblasts (65), but whether galunisertib inhibits phosphorylation of PDGFR, FGFR, and VEGFR in fibroblasts remains unclear. These findings suggest the engagement of these additional receptor tyrosine kinases for the induction of pro-invasive ECM. Activation of PDGFR, FGFR, and VEGFR signaling has been shown to induce a myofibroblastic phenotype in CAFs, leading to enhanced contractile activity and increased deposition of ECM components such as collagen I and fibronectin (66–68). We hypothesize that doxorubicin mediates the release of growth factors upstream of one or more of PDGFR, FGFR, and VEGFR, thereby upregulating signaling pathways beyond TGF-β/Smad3 pathway. Recent study reported a strong correlation between the expression of genes *FAP* and *PDGFA* in human foreskin fibroblasts cultured on patient-derived decellularized tumor scaffolds, showing a link between the expression of the genes associated with CAF marker and ECM remodeling (69). Furthermore, studies using CAFs derived from high-grade serous ovarian cancer patients revealed that CAFs secrete FGF7, which activates autocrine FGF7–FGFR2–PI3K/AKT axis and contributes to maintaining the contractile activity of CAFs (67,70). While our study focused on chemotherapy-driven induction of myCAF-like phenotype in healthy human mammary fibroblasts, recent study demonstrated that doxorubicin increases the expression of *Cxcl9* and *Cxcl10* in mouse cardiac fibroblasts and can promote CD8^+^ T-cell chemotaxis *in vitro* (71).

These chemokines are characteristic markers of interferon-response CAFs, a subtype in which inflammatory and interferon response pathways are highly activated (23). This suggests that doxorubicin can have distinct effects on fibroblasts sourced from different tissues, and also reflects how the plasticity of fibroblasts is largely shaped by their tissue origin (72). Taking these reports and our data together, it is suggested that chemotherapy drugs like doxorubicin can drive changes in the secretome of fibroblasts and engage a broad range of pathways to activate mammary fibroblasts and drive pro-invasive ECM production.

Lastly, our work supports the development of therapies that target the stromal environment to improve the response to standard of care treatment chemotherapy. Our *in vivo* study showed that nintedanib suppressed doxorubicin-induced increases in metastasis, and reduced α-SMA/FAP expression in the primary tumors and the pro-invasive effects of the ECM. Nintedanib is currently FDA-approved for IPF, making it an attractive candidate for drug repurposing, as its established safety profile may facilitate clinical translation. Clinical studies have also supported the therapeutic potential of combining chemotherapy and antifibrotic drugs for the treatment of solid tumors. In the phase III LUME-Lung 1 trial, combination of docetaxel and nintedanib improved progression-free survival compared with docetaxel alone in patients with previously treated non-small cell lung cancer (73). There are also ongoing clinical trials combining chemotherapy and anti-fibrotic drug that target TGF-β signaling in prostate cancer and TNBC (74,75). While our data clearly support the idea that nintedanib is acting in the primary tumor to impact fibroblasts and pro-invasive ECM, it is also possible that it could be acting in the lung metastatic niche to reduce the production of pro-invasive ECM. ECM has been shown to be important for colonization and metastatic outgrowth in the lung (76–78), and chemotherapy has also been reported to remodel the ECM in distant organs. Our previous study has shown that paclitaxel significantly increased the abundance of collagen V in the liver ECM of a TNBC mouse model (79). Paclitaxel has also been reported to drive rapid ECM remodeling in the lung through CD8^+^ T cells expressing lysyl oxidase (LOX), thereby facilitating cancer cell seeding and promoting metastatic colonization (80). Nintedanib exerts potent antifibrotic effects by reducing ECM components such as collagen I in human lung fibroblasts and alleviating overall lung fibrosis in mouse models of pulmonary fibrosis (81,82). Future studies are needed to investigate whether anti-fibrotic drugs can improve response to chemotherapy and reduce recurrence by targeting metastatic sites. Collectively, our study reveals that fibroblasts contribute to the chemotherapy-induced pro-invasive changes in the tumor ECM and highlights the potential of targeting chemotherapy-induced stromal responses as a therapeutic strategy to mitigate the unintended pro-invasive and metastatic effects of chemotherapy.

## Materials and Methods

### *ACTA2* gene expression analysis

Publicly available microarray gene expression datasets containing matched or partially matched breast tumor samples obtained before and after neoadjuvant chemotherapy were downloaded from the Gene Expression Omnibus (GEO). Only patient-matched pre- and post-neoadjuvant chemotherapy samples were included in the analysis. For GSE32603, samples from tumors surgically removed after chemotherapy were used as post-chemotherapy samples. For GSE18728, samples from definitive surgical specimens were used as post-chemotherapy samples. The GEO accession numbers, microarray platforms, numbers of paired samples included in the analysis, and treatment regimens are summarized in Table 1.

### Chemotherapy drugs, inhibitors, and antibodies

Chemotherapy drugs and inhibitors used in this study included doxorubicin (S1208, Selleck Chemicals), paclitaxel (S1150, Selleck Chemicals and J62734.MC, Thermo Scientific Chemicals), cisplatin (S1166, Selleck Chemicals), phosphoramide mustard (HY-137316A, MedChemExpress), nintedanib (HY-50904, MedChemExpress), galunisertib (HY-13226, MedChemExpress), TGF beta-1,2,3 Monoclonal Antibody (1D11) (MA5-23795, Invitrogen), SB202190 (HY-10295, MedChemExpress), and Human DKK-1 Recombinant Protein (120-30-10UG, Gibco). Doxorubicin, paclitaxel, nintedanib, galunisertib, and SB202190 were dissolved in DMSO. Cisplatin was dissolved in DMF. Phosphoramide mustard was dissolved in water. 1D11 and DKK-1 were dissolved in PBS. Primary antibodies included anti-α-SMA, anti-GAPDH, anti-Smad3, anti-phospho-Smad3, anti-Jak2, anti-phospho-Jak2, anti-Stat3, anti-phospho- Stat3, and anti-FAP (Table 2). Alexa Fluor 633-conjugated phalloidin (A22284, Invitrogen) was dissolved in DMSO and used at a 1:500 dilution.

**Table 2.**
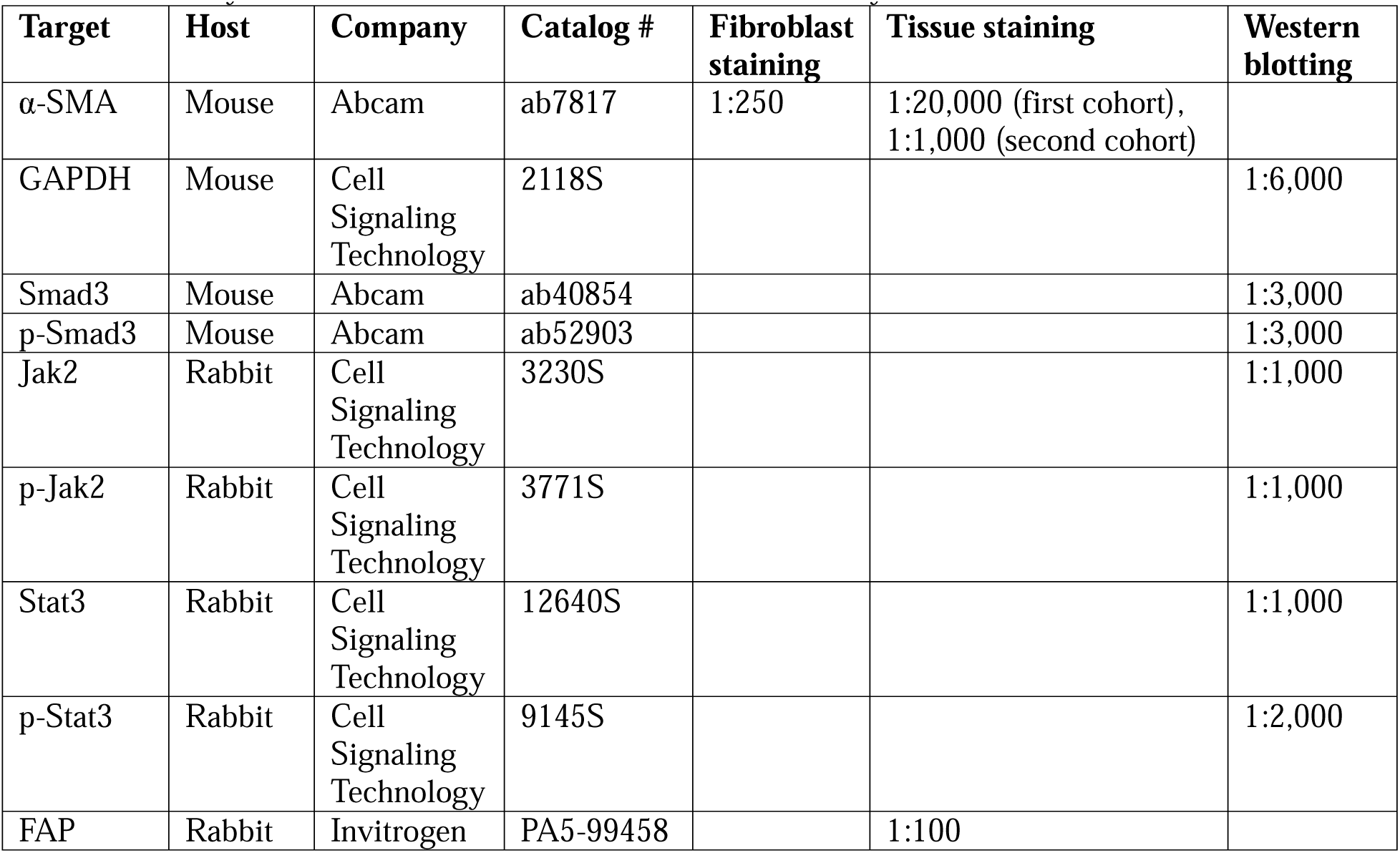
Primary antibodies and dilution factors used in this study.

### Cell culture

MDA-MB-231, MDA-MB-468, SUM159, and EO771 cells were obtained from ATCC. GFP-labeled MDA-MB-231 cells were generated as described previously (20). Reduction mammary fibroblasts (RMFs) (35) and cancer-associated fibroblasts (CAFs) (40) were gifts from Professor Charlotte Kuperwasser’s lab at Tufts University. Cells were cultured at 37°C and 5% CO_2_. MDA-MB-231, MDA-MB-468, and SUM159 cells were cultured in DMEM (10-013-CV, Corning) supplemented with 10% fetal bovine serum (FBS) (1500-500, Seradigm or 97068-085, Avantor) and 1% Penicillin-Streptomycin-Glutamine (PSG) (10378016, Gibco). EO771 cells were cultured in RPMI 1640 medium (11875093, Gibco) supplemented with 10% FBS, 1% PSG, and 10 mM HEPES (15630106, Gibco). RMFs and CAFs were cultured in DMEM supplemented with 10% bovine calf serum (12133C-500ML or 12133C- 1000ML, MilliporeSigma) and 1% PSG. Cells were routinely monitored for mycoplasma contamination using the Universal Mycoplasma Detection Kit (30-1012K, ATCC). Only cells tested negative for mycoplasma were used in this study and all experiments were conducted using the cells at passages up to 20.

### Cell viability assay

MDA-MB-231, MDA-MB-468, or SUM159 were seeded in 96-well plates at 5.0 × 10^3^ cells per well. Cells were allowed to adhere overnight prior to treatment and were then treated with the indicated drug or the corresponding vehicle control for 72 hours. Cell viability was assessed by incubation with PrestoBlue HS Cell Viability Reagents (A13262 ,ThermoFisher) for up to 20 minutes, after which fluorescence was measured at an excitation wavelength of 560 nm and an emission wavelength of 590 nm. Fluorescence from media-only control wells was subtracted from all experimental wells before cell viability was calculated.

### Chemotherapy treatment of fibroblasts for **α**-SMA imaging

RMFs or CAFs were seeded in glass-bottom 24-well plates (P24-1.5H-N, Cellvis) coated with 0.1% gelatin at 500 cells per well for AC-T treatment and 1,000 cells per well for all other treatments.

Fibroblasts were allowed to adhere overnight prior to treatment. For individual drug treatment, RMFs or CAFs were treated with 100 nM doxorubicin, 5 nM paclitaxel, 5 μM cisplatin, or 50 μM phosphoramide mustard, or the corresponding vehicle controls. For AC-T treatment, RMFs were first treated with 50 nM doxorubicin and 50 μM phosphoramide mustard, or the corresponding vehicle controls. Half media changes were performed 24 and 48 hours after the first treatment. 72 hours after the first treatment, a full media was replaced with fresh media and RMFs were incubated for 72 hours. After the 72 hours incubation, half of the media was removed and replaced with fresh media containing 10 nM paclitaxel or vehicle control, resulting in a final paclitaxel concentration of 5 nM. Half media changes were performed 24 and 48 hours after the first paclitaxel treatment. For inhibitor studies, RMFs were treated with 100 nM doxorubicin alone or in combination with an inhibitor, along with the corresponding vehicle controls.

Half of the media were removed and replaced with fresh media containing the experimental condition 24 and 48 hours after the initial treatment. For all treatments, RMFs or CAFs were fixed 24 hours after the last treatment.

### Immunofluorescence staining and confocal imaging of fibroblasts

For immunofluorescence staining of fibroblasts, fibroblasts were permeabilized in 4% PFA, 0.5% Triton X-100 in PBS for 3 min, fixed in 4% PFA in PBS for 20 min, blocked with 10% normal donkey serum (NDS), and incubated with primary antibody (Table 1) overnight at 4°C or 1 hour at room temperature. Fibroblasts were then stained with DAPI and fluorescently labeled secondary antibodies at room temperature for 1 hour. For phalloidin staining, RMFs were incubated with a mixture of Alexa Fluor 633- conjugated phalloidin and DAPI at room temperature for 1 hour. Imaging was performed using Zeiss LSM900 confocal microscope using a 20× objective lens. Multiple adjacent fields of view were acquired and stitched together to generate a single image containing multiple cells.

### Animal experiments

All animal studies were reviewed and approved by the Tufts University Institutional Animal Care and Use Committee. 8-week-old female C57BL/6J mice (000664, The Jackson Laboratory) were injected with 2.0 × 10^5^ EO771 cells suspended in 100 μL of a 20% collagen I (354236, Corning) solution. Tumor size was measured twice weekly using digital calipers in duplicate once tumors became palpable, and tumor volume was calculated using the formula (length × width × width)/2. In the cohort used to compare the vehicle control and doxorubicin groups (Figure 1), treatment with doxorubicin or vehicle control was initiated once the tumors became palpable. In the cohort used for the inhibitor study (Figure 4 and Figure 5), treatment with doxorubicin, nintedanib, or vehicle control was initiated when the average tumor volume reached approximately 50 mm^3^ at 10 weeks of age. Doxorubicin was dissolved in PBS (10010023, Gibco) and administered intraperitoneally at 5 mg/kg every 5 days for a total of four doses in the first cohort. In the second cohort, doxorubicin was administered intravenously at 5 mg/kg once weekly for a total of two doses. Nintedanib was dissolved in 0.5% methylcellulose (HY-125861, MedChemExpress), 0.4% Tween 80 (P1754. Sigma) in 0.9% saline (PB0287, MilliporeSigma) and administered via oral gavage at 50 mg/kg daily on a 5-days-on, and 2-days-off schedule for a total of two cycles. Appropriate vehicle controls were administered to control mice for each treatment. In the first cohort, mice were anesthetized under 3% isoflurane inhalation and euthanized by terminal blood draw and end organ removal under anesthesia 2-3 days after the final doxorubicin administration. In the second cohort, mice were euthanized by CO_2_ inhalation 3 days after the final nintedanib administration. Mice were euthanized earlier if they met institutional humane endpoint criteria, including excessive tumor burden or ulceration. Tumors and lungs were excised and washed in PBS for further study. Tumors were either fixed and embedded in paraffin for H&E and immunofluorescence staining, or transferred into decellularization buffer (0.1% SDS, 1% antibiotic-antimycotic solution). 10% formalin and 4% paraformaldehyde (PFA) in PBS were used as a fixative for the first and the second cohort, respectively. Lungs were fixed in 4% PFA in PBS.

### Hematoxylin and eosin (H&E) staining

Paraffin-embedded tissues were sectioned into 10 μm sections. Sections were deparaffinized, hydrated, and stained with hematoxylin solution Gill No.2 (GHS280-2.5L, Sigma-Aldrich), and Eosin Y solution (HT110180-2.5L, Sigma-Aldrich). Stained sections were mounted in Permount™ Mounting Medium (SP15-100, Fisher Chemical) and imaged using a Keyence BZ-X710 microscope with a 4× objective lens. Lung metastases were manually quantified by two independent observers, and the mean of their measurements was used for analysis.

### Immunofluorescence staining and imaging of tissues

Tumors were sectioned as described above for H&E staining. Sections were deparaffinized, hydrated, and antigen retrieval was performed in Antigen Retrieval Citra Plus Solution (HK080-9K, BioGenex).

Sections were blocked in PBS with 10% normal donkey serum and 0.5% Tween 20 and incubated with primary antibodies overnight at 4°C. Sections were then incubated with secondary antibodies at room temperature for 2 hours and mounted with Fluoromount-G (0100-01, SouthernBiotech). Images were acquired using a Keyence BZ-X710 microscope with a 40× objective lens.

### Reseeding experiments using decellularized tumor ECM

Tumor decellularization and reseeding experiments were performed as described previously (52). Decellularized tumors were cut into 5 mm pieces and pre-conditioned in complete cell culture media overnight. On the day of the experiment, 1.25 × 10^5^ 231-GFP+ cells were reseeded onto the decellularized tumors and allowed to adhere for 6 hours at 37°C. Reseeded tissues were transferred to fresh media, and the cells were imaged for 16 hours, capturing images every 20 minutes using a Keyence BZ-X710 microscope with a 20× objective lens. The cells were tracked using VW-9000 MotionAnalyzer. Cell migration speed, persistence, and mean squared displacement (MSD) were calculated using a custom MATLAB script (MATLAB R2021a, MathWorks). MSD was calculated according to a previous study (83).

### Western blotting

RMFs were seeded in 6-well plates at 1.0 × 10^4^ cells per well for AC-T treatment and at 6.0 × 10^4^ cells per well for all other treatments. RMFs were allowed to adhere overnight prior to treatment. For single-agent treatments, RMFs were treated with 100 nM doxorubicin, 5 nM paclitaxel, 5 μM cisplatin, or 50 μM phosphoramide mustard, or the corresponding vehicle controls, and cell lysates were collected 72 hours after the treatment. For AC-T treatment, RMFs were treated as described above for α-SMA imaging, and cell lysates were collected 24 hours after the final treatment. For inhibitor studies, RMFs were treated with 100 nM doxorubicin alone, an inhibitor alone, the combination of doxorubicin and and inhibitor, or the corresponding vehicle controls. Half media changes were performed 24 and 48 hours after the initial treatment. Cell lysates were collected 24 hours after the final treatment. Proteins were separated by SDS-PAGE using an 8% polyacrylamide gel and transferred to a nitrocellulose membrane using a TransBlot Turbo Transfer System (1704150, Bio-Rad). Membranes were blocked in 5% bovine serum albumin in TBS with 0.05% Tween 20 and incubated in primary antibodies (Table 1) overnight at 4°C with rocking.

Membranes were then incubated with horseradish peroxidase (HRP)-conjugated secondary antibodies at room temperature for 1 hour with rocking. Protein signals were detected using SuperSigna West Femto Maximum Sensitivity Substrate (34095, Thermo Scientific) and imaging was performed using a ChemiDoc MP Imaging System (12003154, Bio-Rad).

### ELISA

RMFs were seeded at 6.0 × 10^4^ cells per well in 6-well plates and allowed to adhere overnight prior to treatment. RMFs were then treated with 100 nM doxorubicin, 5 µM cisplatin, or the corresponding vehicle controls for 72 hours. Culture supernatants were collected and filtered using a 0.22 µm low protein-binding PVDF filter (09-720-3, Fisher Scientific). Cells were fixed and permeabilized using 4% PFA and 0.5% Triton X-100 in PBS as described above, followed by DAPI staining. Stained cells were imaged using a Keyence BZ-X710 microscope with a 10× objective lens. Ten fields of view were acquired per condition, and cell numbers were quantified.

Human TGF-β1 concentrations in culture supernatants were quantified using a commercially available ELISA kit (ELH-TGFb1-1, RayBiotech) following the manufacturer’s protocol. Briefly, culture supernatants were diluted 1:10 in the provided sample dilution buffer. TGF-β1 in the supernatants were activated by the acidification with 1 N HCl, followed by the neutralization with 1.2 N NaOH/0.5 M HEPES. Standards and samples were added to a 96-well plate coated with an anti-human TGF-β1 antibody and incubated to allow antigen binding. Wells were then washed, and a biotinylated anti-human TGF-β1 antibody was added. After washing, HRP-conjugated streptavidin was added, followed by incubation with TMB substrate solution. The reaction was stopped after 30 min using TMB Stop solution, and absorbance at 450 nm was measured using a spectrophotometer. All samples were measured in duplicate, and mean absorbance values were used for calculation of TGF-β1 concentrations. TGF-β1 concentrations were calculated by interpolation from a standard curve generated using GraphPad Prism. To assess TGF-β1 production capacity on a per-cell basis, TGF-β1 concentrations were normalized to the number of cells in the corresponding wells from which the conditioned media were collected.

### Fibroblast-derived ECM generation

Fibroblast-derived ECMs were generated referring to the protocol described previously (84). Briefly, glass-bottom 24-well (P24-1.5H-N, Cellvis) or 48-well plates (P48G-1.5-6-F, Mattek) were coated with 0.2% gelatin for 1 hour at 37°C, followed by cross-linking with 1% glutaraldehyde and quenching of residual aldehydes with 1 M ethanolamine. For reseeding experiments using 24-well plates, RMFs were seeded at 2.0 × 10^5^ cells per well. For experiments using 48-well plates, RMFs or CAFs were seeded at 1.0 × 10^5^ cells per well. For all experiments, fibroblasts were allowed to adhere overnight prior to treatment and were treated with 75 μg/mL L-ascorbic acid together with the indicated chemotherapy drugs or inhibitors to promote robust polymerization and maturation of ECM fibers. For individual drug treatment and inhibitor studies, half of the media were removed and replaced with fresh media containing the mixture of 75 μg/mL L-ascorbic acid and the indicated conditions 24, 48, 72, and 96 hours after the initial treatment to maintain secreted factors while replenishing ascorbic acid and experimental factors. For AC-T treatment, half media changes were performed 24 and 48 hours after the initial treatment. 72 hours after the initial drug treatment, a full media was replaced with fresh media containing 75 μg/mL L-ascorbic acid. Half of the media was removed and replaced with fresh media containing 75 μg/mL L-ascorbic acid 24 and 48 hours after the full media change. Half of the media were removed and replaced with fresh media containing 75 μg/mL L-ascorbic acid and 10 nM paclitaxel or vehicle control, resulting in a final paclitaxel concentration of 5 nM. Half media changes were performed 24 and 48 hours after the paclitaxel treatment. For all treatments, ECMs were decellularized 48 hours after the final half media change.

### Reseeding experiments using fibroblast-derived matrices

ECMs used for reseeding experiments were decellularized following the protocol reported previously (84). After washing the ECMs with DPBS-, ECMs were treated with 20 mM NH_4_OH, 0.5% Triton X-100 in DPBS- for 5 to 15 minutes, until there are no visible cell bodies. After adding the equal volume of DPBS-, the decellularized ECMs were stabilized overnight at 4°C, followed by washing with DPBS- and DPBS+. DNA debris was cleansed using DNAse I (EN0521, Thermo Scientific) at 25 U/mL. 231-GFP+ cells were seeded at 7.5 × 10^3^ cells per well in a 24-well format, and at 1.5 × 10^3^ cells per well in a 48-well format and allowed to adhere to the decellularized ECMs for 6 hours. The cells were imaged for 16 hours, capturing images every 20 minutes using a Keyence BZ-X710 microscope with a 10× objective lens. Data analysis was performed as described above.

### Statistical analysis

Statistical analyses and visualization were performed using GraphPad Prism (version 11.0.1) unless otherwise indicated. For comparisons between two groups, unpaired *t*-test with Welch’s correction, paired *t*-test, or Mann-Whitney test was used. For comparisons of three or more groups, ordinary one-way ANOVA followed by Dunnett’s or Šidák’s multiple comparisons test, mixed-effects analysis followed by Šidák’s multiple comparisons test, or the Kruskal-Wallis test followed by Dunn’s multiple comparisons test was employed. A P-value < 0.05 was considered statistically significant.

### Mean squared displacement (MSD) analysis

MSD was calculated for each individual cell using the following equation:

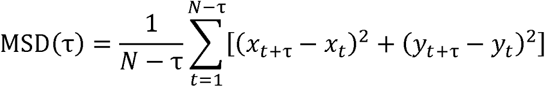

where *N* is the total number of time points (49 in this study), x*_t_* and y*_t_*are the cell position at time *t*, and τ is the displacement interval (1-48 time intervals, corresponding to 20–960 min).

## Author Contributions

**A. Yui:** Conceptualization, data curation, formal analysis, investigation, methodology, project administration, supervision, validation, visualization, writing - original draft preparation, writing - review & editing. **L. Peng:** Data curation, formal analysis, investigation, methodology, visualization, writing - review & editing. **K. C. Lew:** Investigation, visualization, writing – review & editing. **D. J. Landau:** Investigation, visualization, writing – review & editing. **C. Zhang:** Data curation, investigation, methodology, writing – review & editing. **T. J. Gerton:** Investigation, software, writing – review & editing. **M. Li:** Investigation, visualization, writing – review & editing. **S. Jullian Guyard:** Investigation, writing – review & editing. **M. Caron:** Investigation, writing – review & editing. **J. P. Fatherree:** Conceptualization, methodology, writing - review & editing. **R. A. McGinn:** Investigation, writing – review & editing. **A. L. Bayer:** Investigation, methodology, resources, writing - review & editing. **P. Alcaide:** Resources, writing - review & editing. **C. Kuperwasser:** Resources, writing - review & editing.

**M. J. Oudin:** Conceptualization, funding acquisition, project administration, resources, supervision, writing - original draft preparation, writing - review & editing.

## Acknowledgments

This work was supported by NIH (grant no. R01CA255742 and DP2CA271387 to M. J. Oudin, grant no. R01HL144477 and R01HL165725 to P. Alcaide, grant no. F30HL162200 to A. L. Bayer). The authors thank the members of the Oudin lab for their helpful discussions and the staff of Tufts Comparative Medicine Services for animal care. We also thank the Tufts Comparative Medicine Services Histology Core for tissue processing.

---

**Fig. S1.**
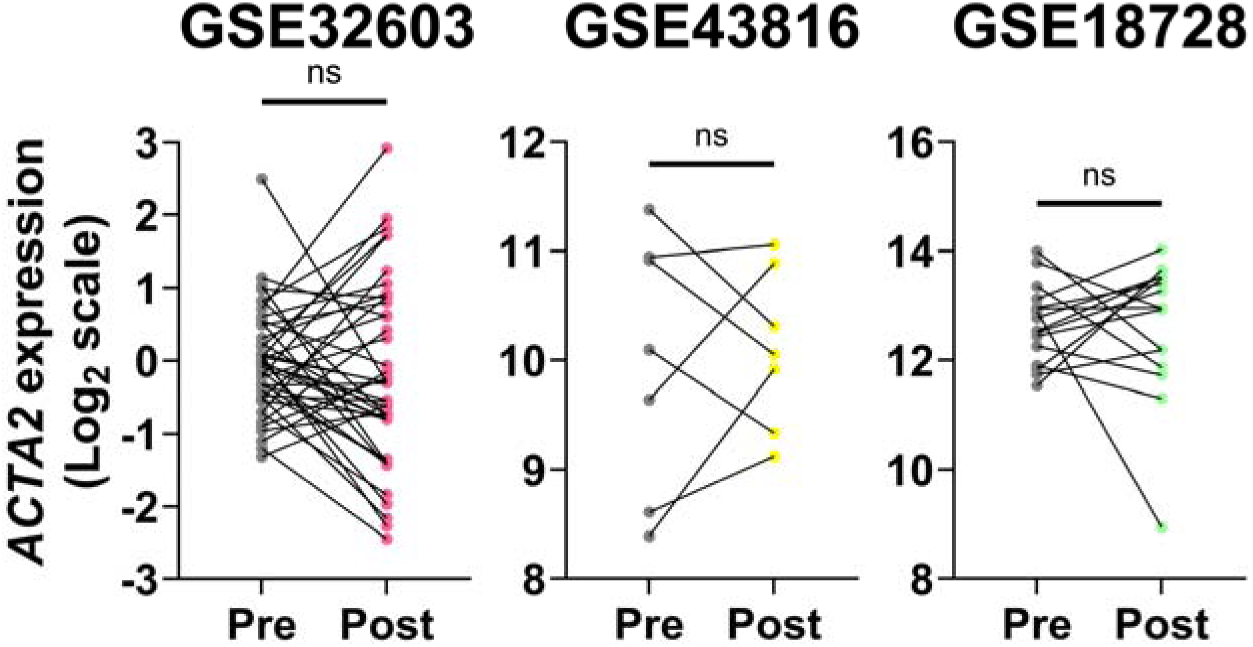
*ACTA2* expression in breast cancer patients pre and post chemotherapy treatment using microarray gene expression datasets. *ACTA2* expression in patient-matched pre- and post-chemotherapy samples. GSE32603: n=43; GSE43816: n=7; GSE18728: n=16 pairs of patient-matched pre- and post-neoadjuvant chemotherapy samples. Significance was determined by paired *t*-test.

**Fig. S2.**
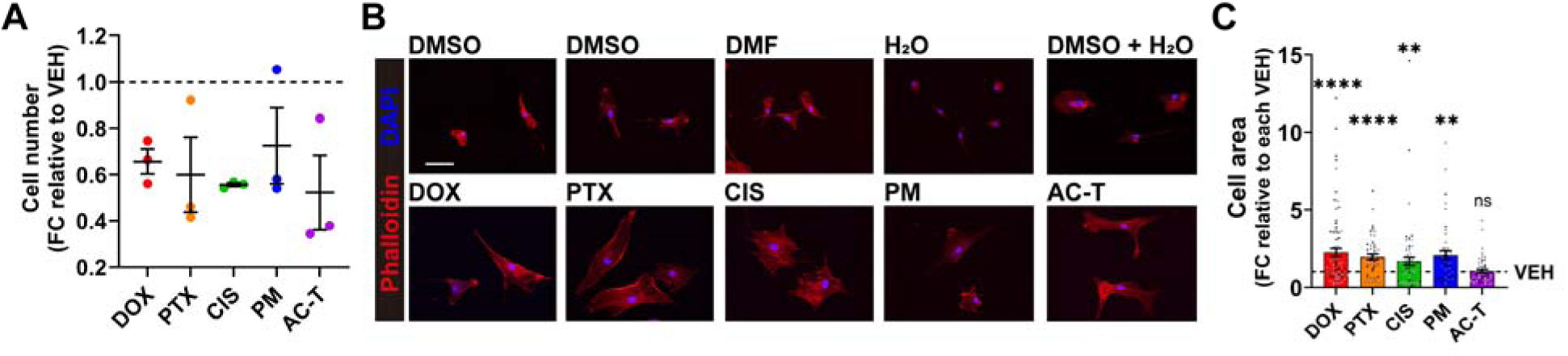
Validation of cell number after chemotherapy treatment and phalloidin-stained RMFs. A) For single-agent treatments, RMFs were treated with 100 nM doxorubicin, 5 nM paclitaxel, 5 µM cisplatin, 50 µM phosphoramide mustard, or vehicle control for 72 h. For AC-T treatment, RMFs were first treated with a mixture of 50 nM doxorubicin and 50 µM phosphoramide mustard or vehicle control for 72 h, followed by a 72 h of incubation in chemotherapy-free media, and subsequent treatment with 5 nM paclitaxel or vehicle control for 72 h. Data are shown as fold change (FC) relative to each vehicle control. B) Representative images of RMFs stained with Alexa Fluor 633-conjugated phalloidin and DAPI. Scale bar: 100 µm. C) Fold change (FC) in cell area was quantified. Data are shown as FC relative to the corresponding vehicle control (VEH). Data show mean ± SEM [n=3 biological replicates, at least 66 cells per condition in total (A), n=2 or 3 biological replicates, at least 44 cells per condition in total (B, C)]. Different-colored data points represent different biological replicates. Significance was determined by unpaired *t*-test with Welch’s correction relative to their own vehicle (C). DOX, doxorubicin; PTX, paclitaxel; CIS, cisplatin; PM, phosphoramide mustard; VEH, vehicle.

**Fig. S3.**
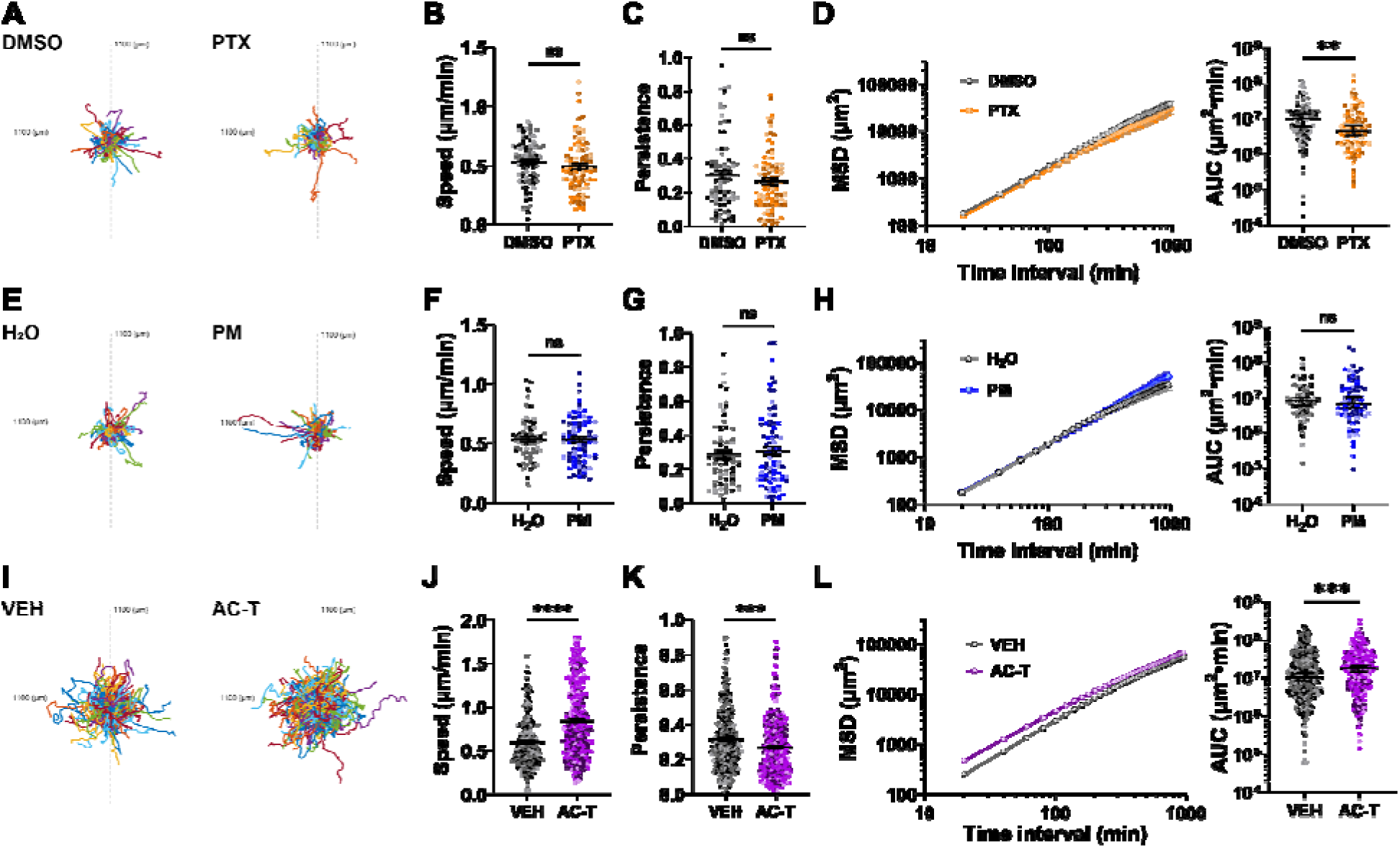
Reseeding experiments using ECM derived from paclitaxel-/phosphoramide mustard-/AC- T-treated RMFs. A, E, I) Trajectories of 231-GFP cells in RMF-derived ECM. The x- and y-axes range from −1100 to +1100 μm. For all treatment regimens, culture media were supplemented with 75 μg/mL ascorbic acid throughout the treatment period. For single-agent treatments, RMFs were treated with 5 nM paclitaxel or 50 μM phosphoramide mustard for 6 days. For AC-T treatment, RMFs were treated with 50 nM doxorubicin + 50 μM phosphoramide mustard for 72 h, followed by a 72 h chemotherapy-free period and then 96 h of 5 nM paclitaxel treatment. B, C, F, G, J, K) Migration speed and persistence of 231-GFP cells reseeded into ECM derived from RMFs treated with paclitaxel (B, C), phosphoramide mustard (F, G), AC-T (J, K), or the corresponding vehicle control. D, H, L) Mean squared displacement (MSD) curves, and MSD AUC of 231-GFP cells reseeded into ECM derived from RMFs treated with paclitaxel (D), phosphoramide mustard (H), or AC-T (L). Data show mean ± SEM (speed, persistence, and MSD curves) or median ± 95% CI (MSD AUC) [n=3 biological replicates, at least 66 cells per condition in total.]. Different-colored data points represent different biological replicates. Significance was determined by unpaired *t*-test with Welch’s correction (speed and persistence), or Mann-Whitney test (AUC) between drug and the corresponding vehicle control. *p<0.05, **p<0.01, ***p<0.001, ****p<0.0001. PTX, paclitaxel; PM, phosphoramide mustard; VEH, vehicle.

**Fig. S4.**
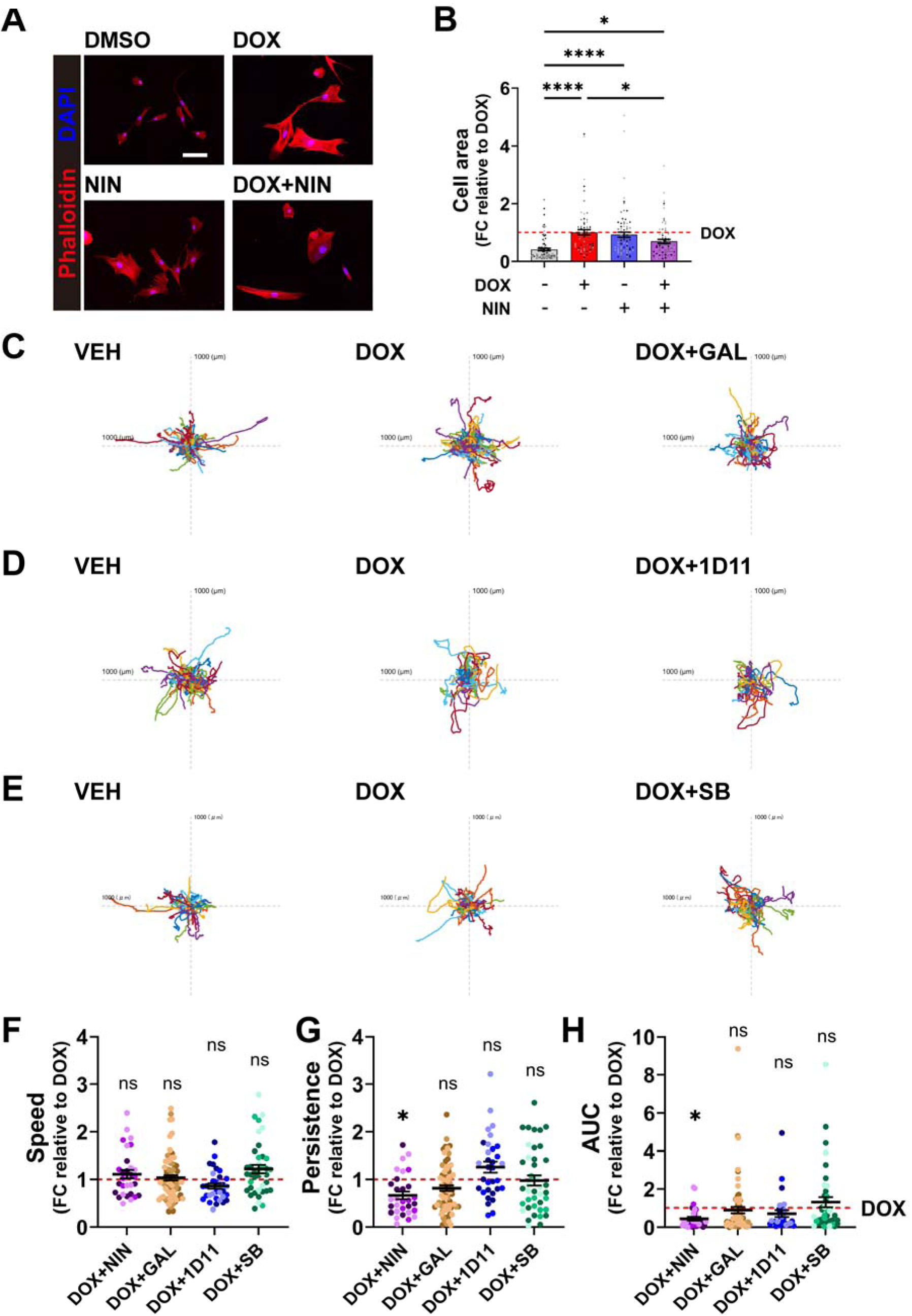
Validation of the effects of inhibitors. A) Representative images of RMFs treated with vehicle control, doxorubicin alone, nintedanib alone, or doxorubicin and nintedanib for 72 h. RMFs were stained with Alexa Fluor 633-conjugated phalloidin and DAPI. Scale bar: 100 µm. B) Fold change (FC) in cell area was quantified. Data are shown as FC relative to doxorubicin. C-E) Trajectories of 231-GFP cells. The x- and y-axes range from −1000 to +1000 μm. RMFs were treated with vehicle control, 100 nM doxorubicin, or 100 nM doxorubicin + one of the inhibitors (1 μM nintedanib, 5 μM galunisertib, 10 μg/mL 1D11 antibody, or 20 μM SB202190) for 6 days in the presence of 75 μg/mL ascorbic acid. F-H) Summary of speed, persistence, and MSD AUC. Data are normalized to the mean value of DOX within each independent experiment. Data show mean ± SEM [n=3 biological replicates, at least 72 cells per condition in total (A, B), or n=3 biological replicates, at least 30 cells per condition in total (C-H).]. Different-colored data points represent different biological replicates. Significance was determined by ordinary one-way ANOVA followed by Šídák’s multiple comparisons test (B) or Dunnett’s multiple comparisons test (F, G), or Kruskal-Wallis test followed by Dunn’s multiple comparisons test (H) comparing DOX with the corresponding vehicle control or DOX + the indicated inhibitor. *p<0.05, **p<0.01, ***p<0.001, ****p<0.0001. DOX, doxorubicin; NIN, nintedanib; GAL, galunisertib, SB, SB202190; VEH, vehicle.

**Fig. S5.**
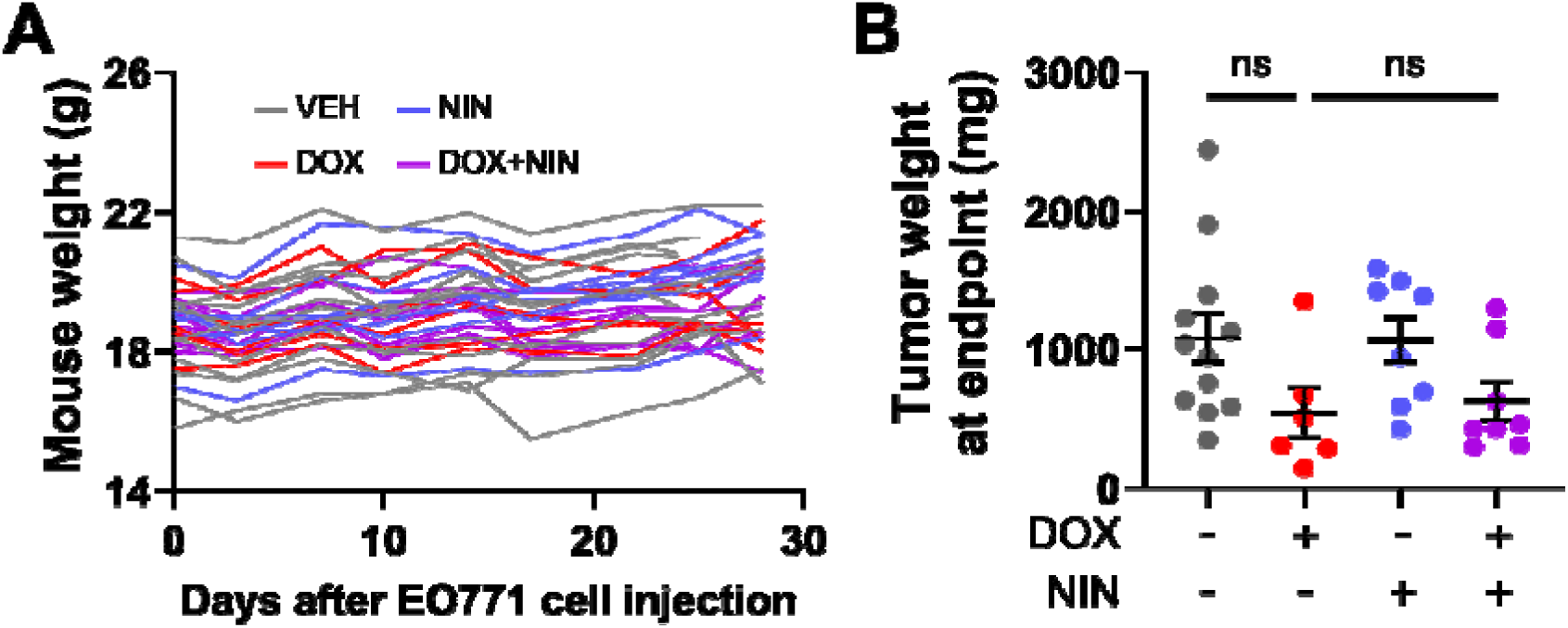
Mouse body weight throughout study and tumor weight at endpoint. A) Mouse body weight was monitored twice weekly as an indicator of treatment-related toxicity. No mice exhibited a body weight loss exceeding 10% of their baseline weight, a threshold that would have prompted immediate veterinary consultation. No mice showed other signs of toxicity, including behavioral abnormalities or loss of body condition. B) Tumor weight at endpoint. Data show mean ± SEM [n=12, 6, 8, and 8 mice for vehicle, doxorubicin, nintedanib, and doxorubicin + nintedanib, respectively.]. Kruskal-Wallis test followed by Dunn’s multiple comparisons test was performed. *p<0.05 was considered statistically significant. DOX, doxorubicin; NIN, nintedanib; VEH, vehicle.

**Supplementary Table 1.**
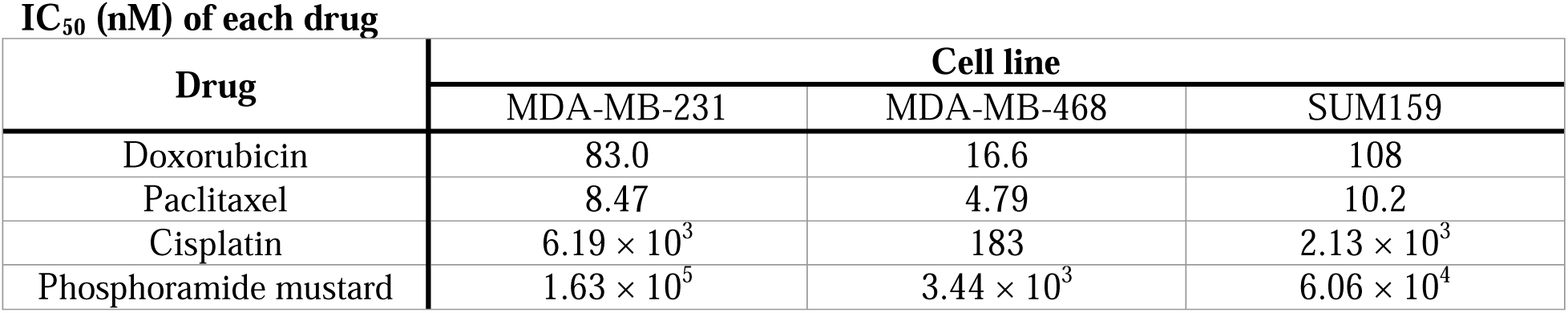
IC_50_ (nM) of each drug Cells were treated with serial dilutions of each drug: MDA-MB-231 (doxorubicin: 0.01-1 μM, paclitaxel: 0.5-25 nM, cisplatin: 1-100 μM, phosphoramide mustard: 1-1000 μM), MDA-MB-468 (doxorubicin: 0.01-1 μM, paclitaxel: 0.5-25 nM, cisplatin: 0.1-10 μM, phosphoramide mustard: 0.1-500 μM), SUM159 (doxorubicin: 0.01-1 μM, paclitaxel: 1-25 nM, cisplatin: 0.1-10 μM, phosphoramide mustard: 0.1-500 μM). Cell viability was normalized to the mean value of vehicle-treated control cells, which was set to 1. Data from three biological replicates, eight (doxorubicin, paclitaxel, cisplatin) or six (phosphoramide mustard) technical replicates in total were pooled, and the mean values were used for analysis. IC_50_ values were determined by nonlinear regression analysis using a variable slope (four-parameter) model.

